# Intracortical bipolar stimulation allows selective activation of neuronal populations in the cortex

**DOI:** 10.1101/2025.03.21.644593

**Authors:** Maarten Schelles, Anke Verhaege, Nils Van Rompaey, Laurens Goyvaerts, Keimpe Wierda, Frederik Ceyssens, Michael Kraft, Vincent Bonin

## Abstract

**Background:** Intracortical electrical stimulation has emerged as a promising approach for sensory restoration, such as a cortical visual prosthesis, yet its effectiveness is limited by current spread and electrode density constraints.

**Objective:** To determine whether intracortical bipolar current steering—via modulation of the return electrode position—can enhance neural activation selectivity compared to traditional monopolar stimulation, with the aim of improving spatial precision in sensory restoration.

**Methods:** We applied intracortical stimulation and used two-photon calcium imaging on acute brain slices to directly visualize neural responses to bipolar stimulation. Biophysical computational modeling was used to complement the experimental results. The analysis included both cellular and population-level assessments to evaluate the impact of several stimulation patterns, such as current direction, electrode spacing and current amplitude, on recruitment patterns.

**Results:** Bipolar stimulation selectively activated distinct neural populations based on the direction of the current flow. This approach decreased the overlap between activated groups and increased the number of independently addressable neural clusters by up to 9-fold relative to monopolar stimulation. Moreover, the electrode configuration and spacing critically influenced the spatial spread of activation.

**Conclusions:** Intracortical bipolar current steering enhances neural activation selectivity by engaging independent neural populations through current directionality. These findings suggest that this strategy may improve the spatial precision of neural prosthetics and sensory restoration without the need for an increased electrode density.

## Introduction

Intracortical electrical stimulation is a promising approach in sensory restoration and brain-computer interfaces [1–6]. Neural prostheses achieve this by using arrays of microelectrodes to deliver small currents, thereby activating neuronal populations and eliciting perceptual or motor responses [7,8]. However, their resolution is constrained by the number of stimulating electrodes, electrode spacing and the spread of electrical fields. Monopolar stimulation, where a single electrode is paired with a distant ground return, produces a direction-independent electric field, often leading to overlapping activation of multiple neural populations [9–12]. This limits the ability to target specific neurons, restricting the spatial precision of neural prostheses. Neuronal responses to electrical stimulation depend not only on the electrode’s location but also on the direction and gradient of the electrical field [13–16]. The orientation of axons relative to the field gradient plays a key role in shaping activation patterns [17–22]. Monopolar stimulation offers limited control over these factors.

Bipolar stimulation can improve spatial selectivity by steering the electric field between two closely spaced electrodes that both actively control the current. This approach allows more precise targeting of neural subpopulations by shaping the electrical field locally [15]. Bipolar stimulation has improved selectivity in cochlear implants [23], deep brain stimulation [22,24,25], spinal cord therapies [26], optic nerve stimulation [27,28], and retinal implants [16,19,29–32]. In cortical visual prostheses, current steering at the cortical surface has enabled the creation of virtual electrodes and more naturalistic phosphene perception [15,33]. Phosphene attributes such as size and brightness are directly influenced by stimulus parameters, including current amplitude [4,34]. Higher current amplitudes recruit larger neural populations, resulting in larger and less precise phosphenes [8,35], highlighting the need for more selective stimulation.

Despite these advances, the effects of bipolar stimulation on activity of neuronal populations in the brain remain poorly understood. Prior studies have primarily focused on surface stimulation or behavioral outcomes, leaving a gap in understanding its influence on neuronal activation patterns. Improving this understanding is essential for enhancing the resolution of visual prostheses, which currently remain constrained by the number of implanted electrodes.

In this study, we used two-photon calcium imaging in acute brain slices and computational modeling to investigate how bipolar stimulation shapes neuronal activation. Using flexible intracortical multi-shank electrode arrays, we analyzed how the direction of return current influences single-cell and population-level activation, focusing on current amplitude, pulse polarity, waveform asymmetry and electrode spacing. Additionally, we employed a biophysically realistic simulation framework to complement experimental findings. Our results show that bipolar stimulation enables more selective activation of neural populations by steering current flow, demonstrating that neuronal recruitment depends on both electrode placement and current directionality. By systematically characterizing how current steering shapes neural activation, this study provides insights for enhancing resolution in neural prosthetics beyond the physical constraints of electrode density.

## Materials and Methods

### Animals and slice preparation

All experiments were approved by the local ethical committee on animal experiments of KU Leuven. We used CaMkII-tTA x TREGCaMP6s mice [59] which have efficient expression of GCaMP6 in the primary visual cortex region (n = 11, aged 30 to 90 days, mixed male and female). Mice were anesthetized using 2ml of isoflurane and after decapitation, the brain was quickly removed and transferred into ice-cold cutting solution (in mM): 87 NaCl, 10 glucose, 75 sucrose, 2.5 KCl, 1.25 NaH2PO4, 0.66 CaCl2, 15 MgCl2, 5 ascorbic acid, 10 kynurenic acid, 22.5 NaHCO3, 3 pyruvic acid, saturated with 95% O2 / 5% CO2 [60,61], and whole brain coronal slices (300 µm) containing primary visual cortex were cut using a vibratome (VT1200, Leica Biosystems) [62–64]. The neural tissue was sliced and, after a recovery phase in a 34°C water bath, stored at room temperature in cutting solution. During GCaMP6 imaging experiments slices were continuously perfused in a submerged chamber (Warner Instruments) at a rate of 3–4 ml/min with aCSF (in mM): 2 CaCl2, 127 NaCl, 2.5 KCl, 1.25 NaH2PO4, 1 MgCl2, 25 NaHCO3, 25 D-glucose, saturated with 95% O2 / 5% CO2 [65] (Fig. 1C).

**Figure 1:**
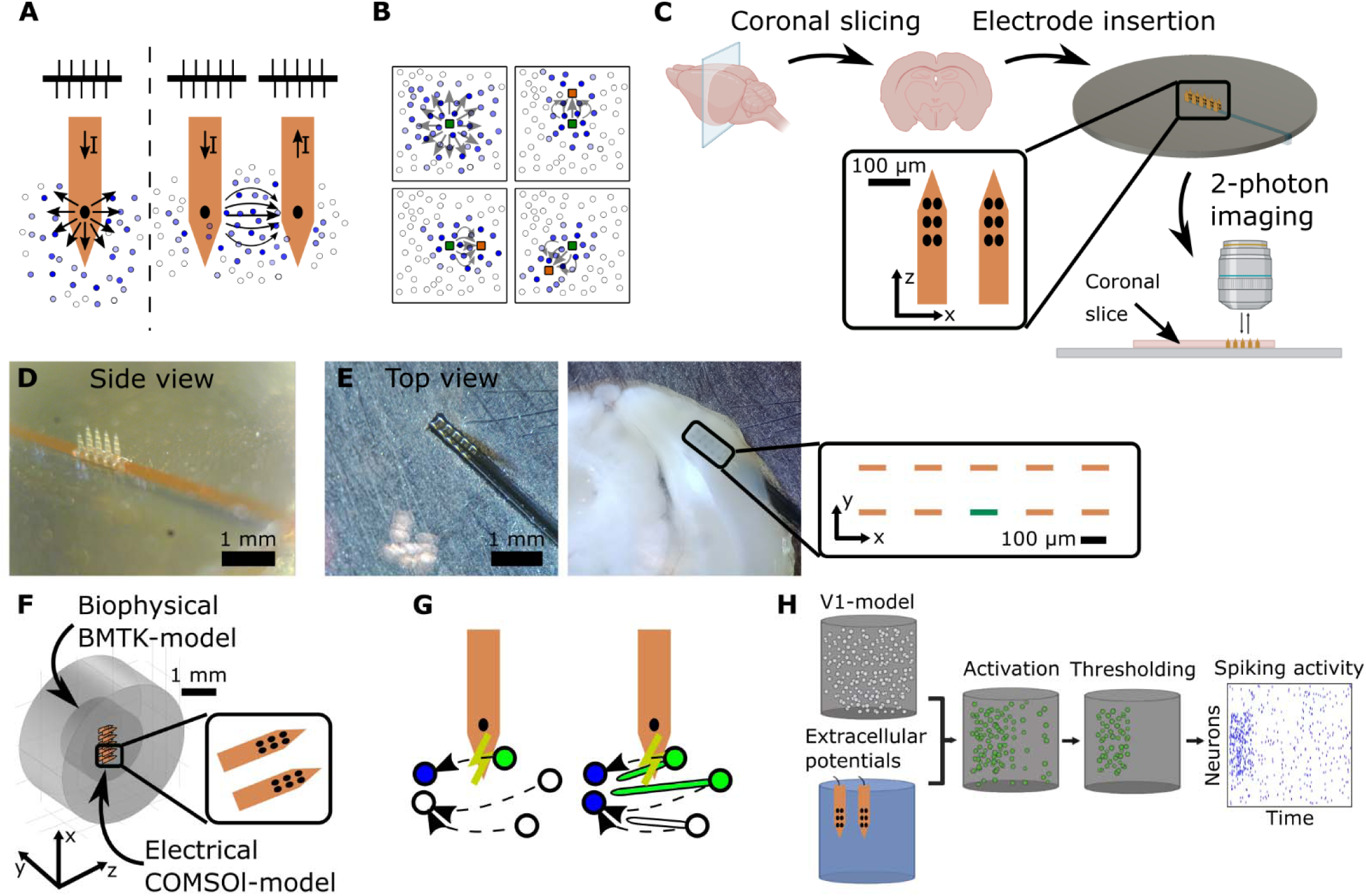
Two-photon calcium imaging experiments and simulation environment to investigate the neural response to bipolar stimulation. **(A)** Monopolar (left) vs. bipolar stimulation (right). **(B)** Electrical charge spreads direction-independent, or spherically, for monopolar stimulation (top left) and direction-dependent for bipolar stimulation, influenced by return electrode placement, with the green and orange squares defined as fixed central and variable return electrode, respectively. Activated neurons are indicated as blue circles, with darker shades representing higher activity levels. Using bipolar stimulation, the activity can selectively be shifted around the green electrode, compared to monopolar stimulation. **(C)** Experimental procedure: coronal slicing, electrode insertion, and calcium imaging using custom flexible electrodes. Imaging performed 100 µm below the slice surface, aligning with the top electrode row. **(D)** Electrode array: two rows of five flexible shanks, each with six electrodes. **(E)** Top view of the electrode array before and after insertion in the coronal slice. In the enlarged view, the green rectangle denotes the fixed central electrode. **(F)** COMSOL model: biophysically realistic inner cylinder, including a model of the electrode arrays, and an outer cylinder using a leaky-integrate-and-fire model. **(G)** Incorporating axons in the biophysical model improved simulated activation of distant neurons through extracellular stimulation. Green circles represent neurons activated by the electrical stimulation, blue circles represent indirectly activated neurons, via synaptic transmission. **(H)** Thresholding was performed on simulated neural activity after extracellular stimulation, accurately determining the cellular activation caused by extracellular stimulation.

### Electrophysiological setup and electrode design

Custom electrode arrays were fabricated with five polyimide shanks, transparent under two-photon imaging (150 µm pitch, 20 µm thick, 400 µm long) [66,67]. Each shank had three rows of two iridium oxide electrodes (30x20 µm², 15 µm apart, 31 ± 10 kΩ impedance, measured at 1 kHz). Two arrays were mounted on a stainless steel backplate (150 µm apart), with shanks up and electrodes facing each other (Fig. 1D, E). No distant large return electrode was used, the chamber remained floating. Electrode insertion was performed in sucrose solution, with the slice positioned using pipettes and lowered onto electrodes, with the top array in L2/3, and bottom array in L4. The setup was transferred to aCSF in a two-photon chamber.

### Electrical stimulation

Electrical stimulation was applied using a custom neural stimulator (0.7 µA resolution). Switch arrays allowed independent, timed bipolar stimulation of electrodes, controlled by a custom Python script. Synchronization with the recording setup used a TTL-pulse, and electrode voltages were monitored with a PicoScope 2000 (Pico Technology).

Pulse trains consisted of 40 charge-balanced biphasic pulses (200 µs width, 200 Hz). Pulses were symmetrical or asymmetrical (second phase four times longer and lower amplitude). Stimulation was bipolar between a central and one or two return electrodes, with opposite currents. Polarity was defined as the leading phase of the central electrode. Electrodes were shorted between pulses to ensure charge balance, and unused electrodes remained high-impedant.

### Two-photon calcium imaging

The aCSF temperature in the recording chamber was maintained at 34 ± 0.2°C using a temperature controller (Luigs & Neumann). Imaging was conducted using fast-gated GaAsP PMTs (Hamamatsu H11706) with a two-photon microscope (Vivo Multiphoton, 3i) and a Mai Tai HP laser (920 nm excitation, Spectra-Physics). A 16x water immersion objective (N16XLWD-PF, Nikon) captured 9 Hz frames with a 505x485 µm² field of view (397x380 pixels) at 100 µm depth, aligning with the top row of microelectrodes and minimizing damaged neurons at the surface. Each recording applied 4–8 stimulation patterns, repeated 6 times, interleaved with the other patterns, and spaced 8 seconds apart to ensure independent calcium responses [68].

### Simulation framework

The in-silico model used is a hybrid approach with two main steps [69]. First, extracellular potentials from electrical stimulation were calculated using a Finite Element Model (FEM) in COMSOL, designed to match the geometry and configuration of the calcium imaging setup. These potentials were applied to an existing biophysically realistic neural network model [36] simulated with the Brain Modeling Toolkit (BMTK) and NEURON [37,70]. Based on the mouse visual cortex, the model includes a cylindrical core (800 µm height, 400 µm radius) of biophysical V1 neurons surrounded by an annular region with leaky integrate-and-fire neurons, representing lateral geniculate nucleus and background neurons (845 µm total radius). The core is divided into five layers, combining layers 2/3 due to similar properties (Fig. 1F).

The biophysical models simulate neurons as electrical circuits with detailed morphologies and dynamics. To improve the accuracy of extracellular stimulation, we added active axons to the model, replacing passive axon stubs with alternating segments containing nodes of Ranvier and myelinated sections (Fig. 1G). The passive sections maintained a high-resistance, low-capacitance membrane to mimic the behavior of myelinated axons (Table 1). Active compartments included sodium and potassium channels [71], modeled using NMODL and integrated into BMTK/NEURON. Computational resources were provided by the Flemish Supercomputer Center (VSC).

**Table 1:** Electrical axonal properties in the biophysical model.

| Model Parameters | Value |
| --- | --- |
| Length compartment active | 30µm |
| Length compartment myelin | 30µm |
| Diameter compartment active axon | 1µm |
| Diameter compartment myelin | 1µm |
| Ra | 150 Ω cm |
| ENa | 55 mV |
| EK | -77 mV |
| EL | -70 mV |
| C<sub>active</sub> | 1 µF/cm<sup>2</sup> |
| g<sub>Nat</sub> | 1,5 S/cm<sup>2</sup> |
| g<sub>Nap</sub> | 0,002 S/cm<sup>2</sup> |
| g<sub>K</sub> | 1,6 S/cm<sup>2</sup> |
| g<sub>I,active</sub> | 0,04 S/cm<sup>2</sup> |
| C<sub>myelin</sub> | 0,005 µF/cm<sup>2</sup> |
| g<sub>I,myelin</sub> | 0,000 0015 S/cm<sup>2</sup> |

### Data processing and statistical analysis

Raw recordings were processed in ImageJ. When needed, recordings were corrected for drift in the x- and y-directions. The fluorescence response (ΔF/F) was calculated as the difference between fluorescence in the 8 seconds post-stimulation and the baseline (3 seconds pre-stimulation), normalized to the baseline, providing a temporal signal per pixel. To enhance signal-to-noise-ratio, signals were averaged across 6 repetitions of the same stimulation pattern within a recording. Variance was assessed by comparing the first 3 repetitions with the latter 3. Morphological segmentation using Cellpose 3 identified regions of interest (ROIs) as cell bodies with visible ΔF/F after stimulation [72,73].

Data analysis was conducted using custom Python scripts. Calcium traces of each cell were calculated by averaging all pixels within an ROI. The peak fluorescence in the first second post-stimulation was defined as the cell’s response to a stimulation pattern. A cell was considered active if this peak exceeded the mean plus three times the standard deviation of its fluorescence during the 3 seconds before stimulation onset. The cell centroid was determined as the mean position of its pixels in the x- and y-directions.

Similarly in simulations, a cell was defined as active if its spike rate exceeded the mean plus 3 times the standard deviation (Fig. 1H). While the simulation included all cortical layers, actual data analysis only included cells in 2/3 and L4 for direct comparison with calcium imaging data.

To compare two stimulation patterns at the cellular level within the same slice or recording, two metrics were introduced. Binary overlap was the percentage of cells active for both patterns relative to the total cells active for at least one pattern. The Spearman correlation coefficient was defined as 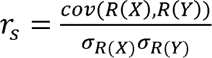 with cov() and σ the covariance matrix and standard deviations, respectively, and R(X) and R(Y) the ranks of X and Y. This was computed using the fluorescence of all cells that were active for at least one stimulation pattern in this particular recording (not necessarily one of the two patterns under comparison).

The angle between two stimulation directions was calculated as the angle between the vectors defined by the central electrode and the return electrodes. For return to two electrodes, the vector pointed to the midpoint of the two.

To compare all different stimulation patterns on a cellular level, we applied principal component analysis (PCA) on the activity of all cells that were active for at least one stimulation pattern, in an n-dimensional space, with n the number of stimulation patterns. Cumulative variance was computed, using the principal components in ascending order.

To compare the centers of neural activity for different patterns, cells were projected onto a main axis, such as the axis defined by the central and return electrode(s) or the cortical layer axis (parallel to the brain surface). A Gaussian fit, weighted by cell fluorescence, was applied to their positions along this axis, yielding the activity center and extent. If too few neurons were available for a Gaussian fit, the mean and standard deviation of the projected neurons were used instead.

## Results

We investigated whether bipolar stimulation increases the selectivity of electrical stimulation and improves resolution for next-generation visual prostheses, compared to monopolar stimulation (Fig. 1 A,B). Using two-photon imaging in acute coronal slices from GCaMP6s mice, we recorded neuronal calcium responses to stimulation through a flexible intracortical multi-shank electrode array (Fig. 1 C-E) and examined how changes in current direction influence neural activity patterns. Neural activity was recorded at 100 µm below the slice surface with a 505 × 485 µm² field of view, focusing on neurons in layers 2/3 and 4. Using data from 11 animals (1–2 slices per animal), calcium responses were quantified over 8 seconds following stimulus onset.

To complement the experimental results, we performed computational simulations using BMTK to model neuronal responses to bipolar stimulation, based on the cellular composition of the mouse visual cortex [36,37]. We used COMSOL to simulate the electrical field perturbation in response to bipolar stimulation (Fig. 1F) and examined the resulting spiking responses in BMTK (Fig. 1 G-H).

To approximate stimulation conditions relevant for visual prostheses, we applied 200 Hz pulse trains lasting 200 ms [4,8,38]. Stimulation amplitudes ranged from 6 to 36 µA, with most data acquired at 10 and 20 µA. We systematically examined how bipolar electrode configuration, electrode spacing, amplitude, and waveform polarity and asymmetry affected the recruitment and spatial distribution of activated neurons.

### Cells and neural populations are activated preferentially by different stimulation directions

We examined how bipolar stimulation through different electrode configurations influences neuronal activation patterns (Fig. 2A). Stimulation was applied with 13 different bipolar electrode pairs, each consisting of a fixed, central stimulation electrode combined with one or more variably positioned return electrodes within the imaging plane (Fig. 2B-D).

**Figure 2:**
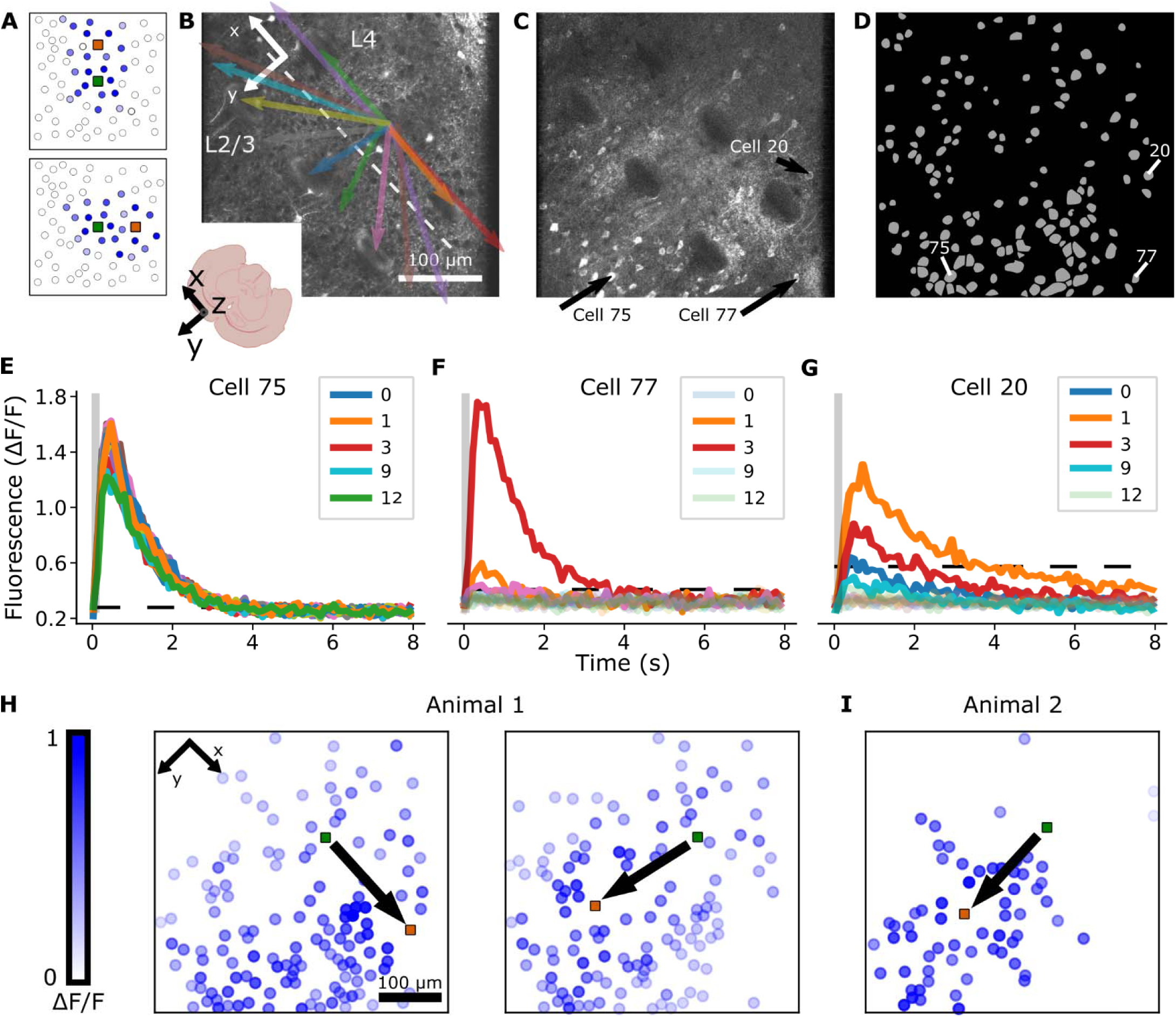
The direction of bipolar stimulation shapes the spatiotemporal responses of cells. **(A)** Schematic of the bipolar electrode configuration consisting of a fixed stimulation electrode (green square) and variable return electrode (orange square) showing how stimulation direction influences neuronal activation (blue circles, darker shades representing higher activity). **(B)** Coronal view of the electrode placement in a brain slice, illustrating 13 different bipolar electrode pairs with their corresponding directions. A fixed central electrode in layer 4 (L4) serves as the stimulation source, while 9 variable return electrodes and 4 additional return electrode (combinations of 2 adjacent bottom-row electrodes with 50% of the current each) complete the configuration and act as current sink. The dashed white line represents the border between L2/3 and L4. The white axes indicate the orientation of the coordinate system as defined in the coronal slice. **(C)** Overlay of ΔF/F responses for all 13 bipolar electrode pairs, highlighting the spatiotemporal dynamics of neuronal activation. **(D)** Identification of regions of interest (ROIs) based on peak ΔF/F values during stimulation. (E-G) Temporal fluorescence (ΔF/F) responses of individual neurons to 13 bipolar electrode pairs. Three example cells are shown with different response properties: **(E)** consistently active across all electrode pairs, **(F)** selectively active for a single electrode pair, or **(G)** active for multiple electrode pairs but showing a preference for specific ones. The grey box indicates the onset and duration of stimulation. The colour scheme corresponds to the electrode pairs shown in panel B, with the opaque and transparent lines representing the cell’s activity and inactivity for this bipolar pair, respectively. **(H)** Representative sample of the spatial distribution of ΔF/F activity for two electrode pairs (indicated by coloured squares and black arrows) revealed distinct but overlapping neural populations, with one current direction evoking stronger activity in certain cells. ΔF/F was normalized between 0 and 1 for visualization purposes, with dark blue cells showing the highest ΔF/F for this stimulation, and white cells showing no activity. **(I)** Similar activation patterns were observed for the same stimulation direction in a second animal, demonstrating consistency across experiments.

Neurons exhibited selective responses to different bipolar electrode configurations. Calcium peaks occurred within 1 second of stimulation onset, followed by a slow decay. Some neurons were consistently activated across all bipolar pairs, likely due to proximity of their axon or axon initial segment to the central electrode (Fig. 2E) [9,39,40]. In contrast, other neurons exhibited strong directional preferences, responding preferentially to one (Fig. 2F) or several bipolar pairs (Fig. 2G), likely influenced by the position of the return electrode.

To assess activation at the population level, we identified neurons with significant responses to stimulation (>3 times the standard deviation of the pre-stimulus fluorescence) and examined their peak fluorescence (ΔF/F). Different electrode pairs produced distinct spatial activation patterns (Fig. 2H). Although the precise location of activated neurons varied, the overall population response patterns were similar across the recordings in different slices (Fig. 2I and Fig S1). These observations suggest that intracortical bipolar stimulation can target distinct neuronal populations by adjusting the direction of the stimulation current.

### A biophysical model with axons responds more realistically to extracellular stimulation

To examine how specific activation patterns emerge from bipolar stimulation, we simulated neuronal responses using a biophysical model. Incorporating axons into the model enabled activation of cell bodies hundreds of micrometers away from the bipolar pair (Fig. 3A, B, Table 2), aligning with experimental observations of extracellular stimulation [9,39,41]. While the model included all cortical layers, only activity in L2/3 and L4 was visualized and analyzed further.

**Figure 3:**
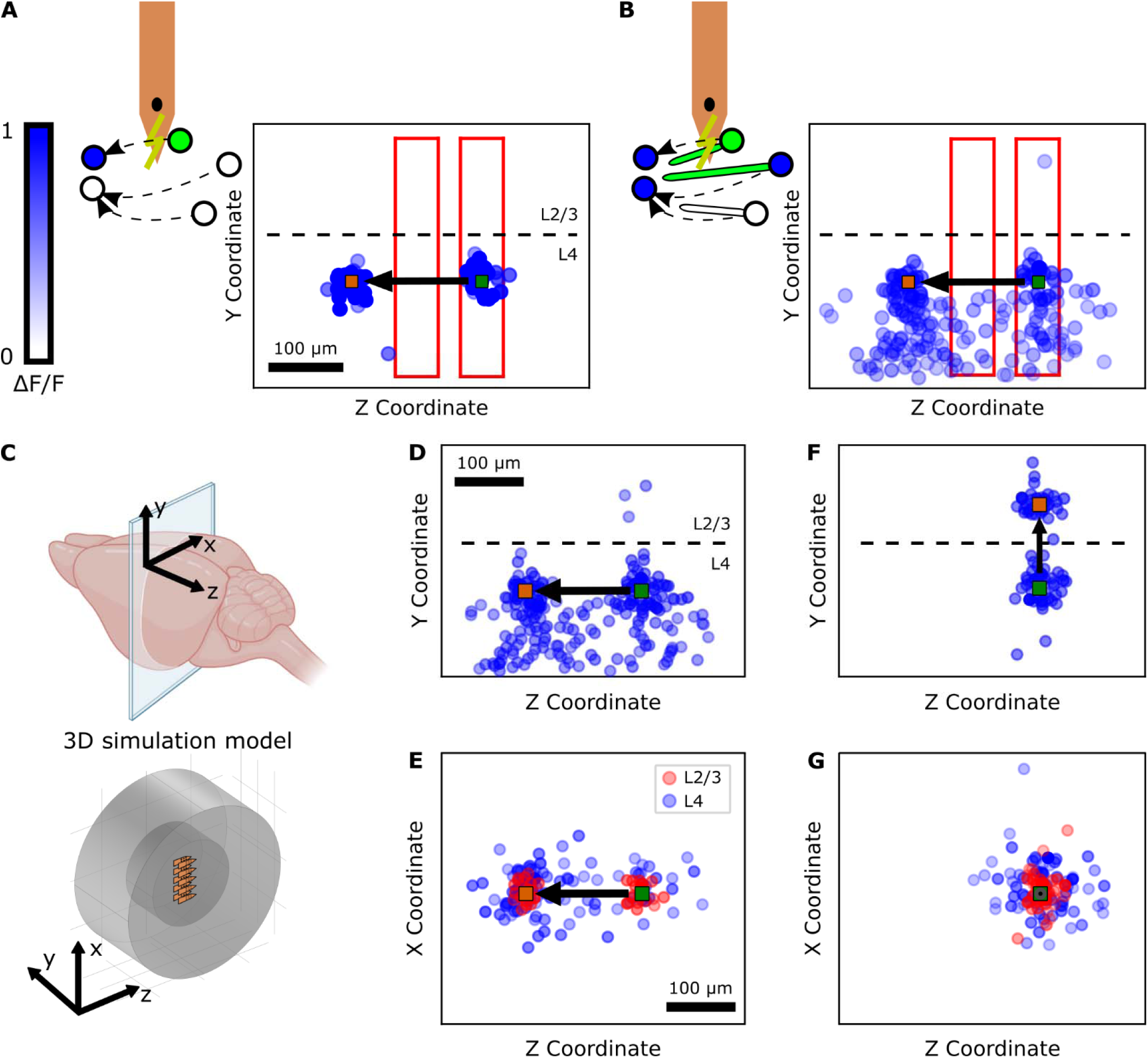
Inclusion of axons in the simulation model increases the realistic effect of stimulation. **(A)** Simulated neuronal activation in the model without axons, and **(B)** the updated model incorporating axons (10 µA stimulation along cortical layer). Dashed lines indicate the boundary between cortical layers 2/3 and 4. Blue circles represent activated neurons, with colour intensity indicating a higher ΔF/F. Green and orange squares denote the fixed central and variable return electrode positions, respectively. Red rectangles define the regions used for neuron and spike counts (Table 2). **(C)** A 3D computational model provided coronal and horizontal views of the simulated cortical response. The coordinate system was defined in the same way as in the coronal slices (Figure 2) **(D, E)** Activation patterns when current (20 µA) was applied along the cortical layer, shown in (D) coronal and (E) horizontal views. In the horizontal view, neurons in L2/3 and L4 are coloured red and blue, respectively, for visualization purposes. **(F, G)** Activation patterns when current (20 µA) was applied along the cortical column from L4 to L2/3, shown in (F) coronal and (G) horizontal views.

**Table 2:** Comparison of neuronal activation patterns in response to bipolar stimulation, with and without axons.

| Model | Location | Number of active neurons | Number of spikes |
| --- | --- | --- | --- |
| Model excluding axons | Around electrodes | 22 | 422 |
|  | In between electrodes | 0 | 0 |
| Model including axons | Around electrodes | 35 | 634 |
|  | In between electrodes | 13 | 170 |

The model provided both coronal and horizontal views of neural activation (Fig. 3D-G). Similar to the imaging experiments, different bipolar electrode pairs selectively activated distinct neuronal populations (Fig. 3D,F and Fig. 3E,G).

### Decreasing direction similarity reduces neural population overlap

To quantify the similarity of populations activated by different stimulation directions, we measured the overlap of active neurons across bipolar electrode pairs in both experimental and simulated data. Overlap was defined as the proportion of neurons active for both electrode pairs relative to those active for either pair. Additionally, we computed the Spearman correlation coefficient between the peak response amplitudes.

We compared all combinations of electrode pairs within a single recording that shared a fixed central electrode but had different return electrodes. Some electrode pair combinations exhibited high overlap, where similar neurons were activated by both stimulation conditions, yielding a high correlation coefficient (Fig. 4A, B). In contrast, other combinations had low overlap, with certain neurons inactive or weakly activated for one of the two pairs (Fig. 4C-F). In these cases, correlation coefficients were less meaningful (Fig. 4F), so overlap was used as the primary metric for assessing population similarity.

**Figure 4:**
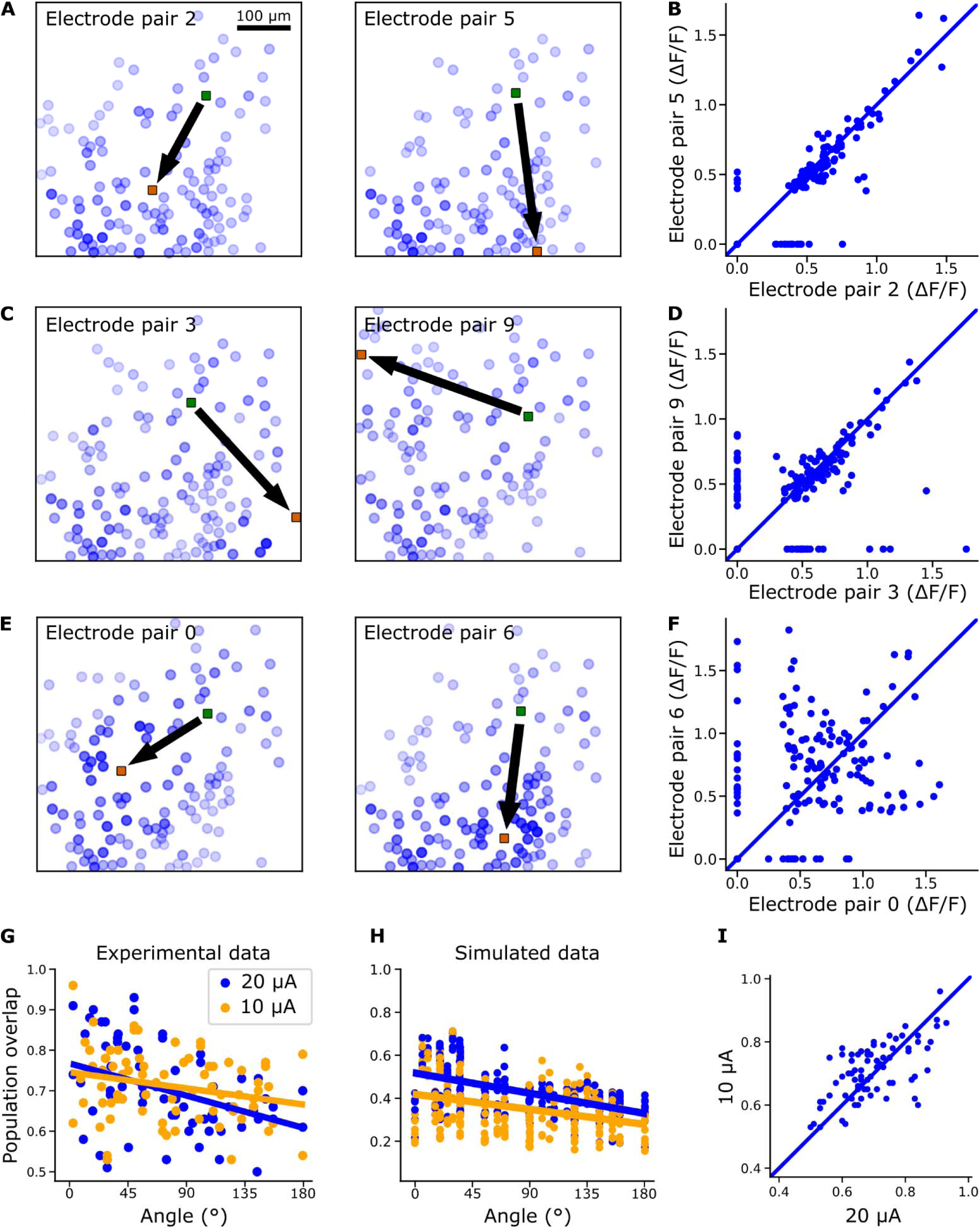
Different current directions (same central electrode, variable return electrode) result in partially overlapping neural activation. **(A)** Spatial activity maps for bipolar electrode pairs with a small enclosed angle show high overlap, with most neurons being co-active and exhibiting similar fluorescence responses (ΔF/F), represented by a similar shade of blue (darker blue means higher activity). The fixed central and variable return electrodes (green and orange squares) and current direction (black arrow) are indicated. **(B)** Pairwise correlation of ΔF/F between these conditions is high (r = 0.88), the identity line is shown as comparison. **(C, D)** For less similar electrode pairs, some neurons remain co-active with comparable ΔF/F, while others are either inactive or only active for one condition, leading to a moderate correlation (r = 0.47). **(E, F)** When electrode pairs are highly different, neurons show minimal co-activation, and ΔF/F responses vary significantly, resulting in a low correlation (r = 0.22). **(G)** The overlap between neural populations decreases as the enclosed angle between electrode pairs increases at both 10 µA and 20 µA stimulation intensities (blue: 20 µA, orange: 10 µA). The full line represents the linear fit for both intensities. **(H)** A similar trend was observed in silico. **(I)** Across experiments, neural population overlap in function of the angle was moderately correlated for 10 µA and 20 µA when measured in the same slice (paired cellular data, r = 0.58).

The overlap between neural populations decreased as the angle between electrode pairs increased, from 0° to 180° (Fig. 4G). Linear fits revealed overlaps of 74% and 76% at 0° for 10 and 20 µA, respectively, with slopes of -0.04% and - 0.09% per degree. In simulations (Fig. 4H), overlaps were lower (42% and 52% at 0°) with slopes of -0.08% and -0.10%. While both experimental and simulated data exhibited a similar trend, the higher overlap in experiments is likely due to the planar focus of calcium imaging, which captures a limited subset of the 3D activation volume modeled in simulations.

Similar results were obtained across current amplitudes. A paired analysis of response overlaps at 10 µA and 20 µA for the same electrode pairs showed a high correlation (r = 0.58, correlation between 10 µA and 20 µA) (Fig. 4I), suggesting that direction-dependent activation patterns remain stable across stimulation intensities.

### A number of independent neural populations exist when sending current into different directions

As the angles between two stimulation directions decreased, the overlap of the neural activity increased, indicating a limit to the number of distinct neural populations that can be activated within a given area. This suggests that the number of distinct neural populations is lower than the number of stimulation directions, indicating a reduced dimensionality of neural activation patterns.

To assess this dimensionality, we applied principal component analysis (PCA) to the peak response of each neuron after stimulation, using both experimental and simulated data.

PCA revealed 4 to 6 distinct neural activation patterns. At a 90% variance threshold, the number of patterns was 4.1 ± 1.1 for 20 µA and 4.0 ± 0.8 for 10 µA, while at a 95% threshold, the number increased to 6.0 ± 1.3 for 20 µA and 6.0 ± 1.0 for 10 µA (Fig. 5A, B). These results demonstrate a high consistency of the number of distinct neural activation patterns across different recordings and different amplitudes.

**Figure 5:**
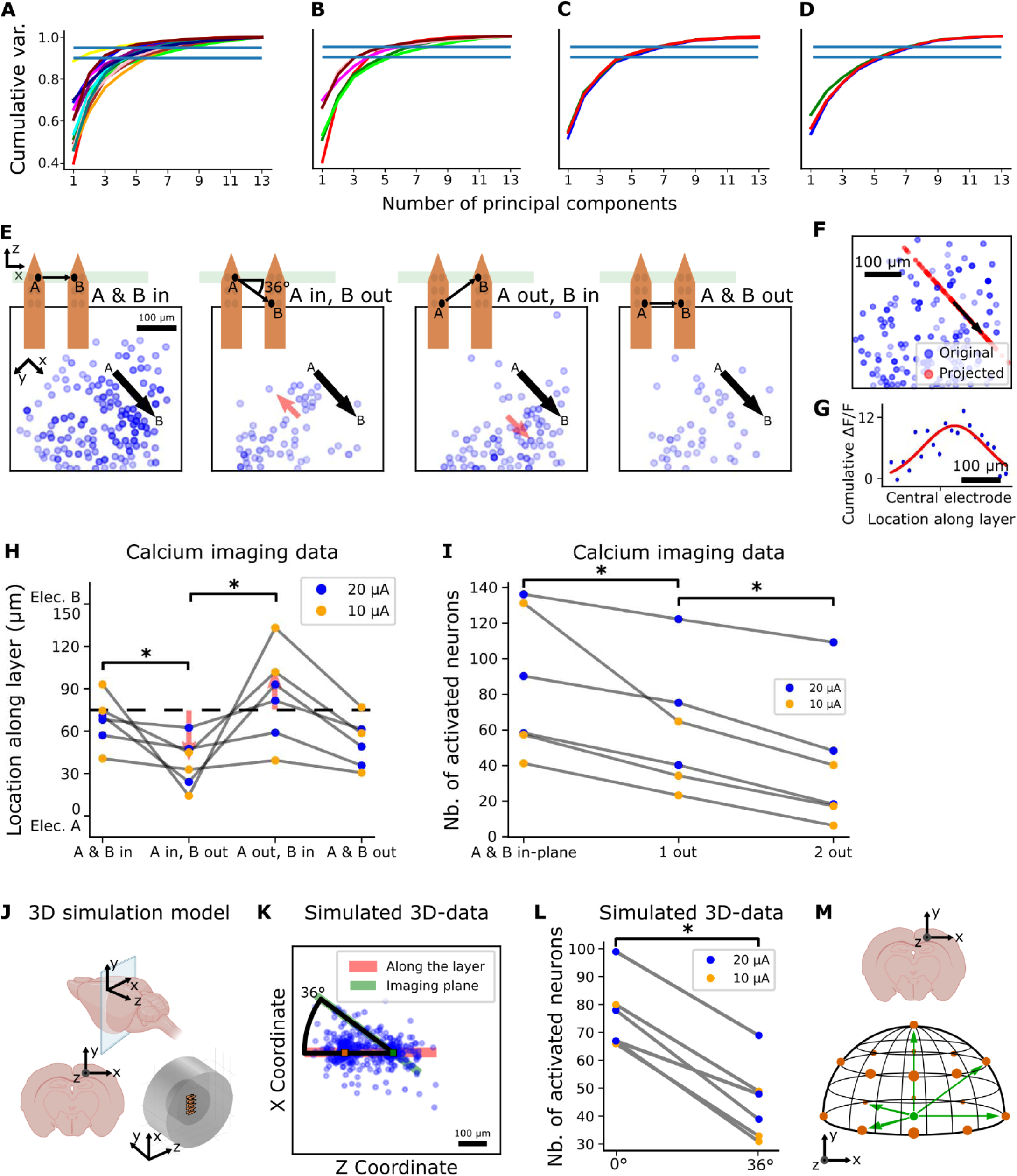
Sending current in different directions yields lower-dimensional activation patterns. PCA of cellular activity for 13 different bipolar electrode pairs revealed a lower dimension of independent neural populations in calcium imaging data with **(A)** 20 µA and **(B)** 10 µA, and simulations with **(C)** 20 µA and **(D)** 10 µA. The same colors were used for recordings in the same animal. The blue dashed lines represent 90% and 95% cumulative variance. **(E)** Current sent to or from an electrode outside the imaging plane activated different neural populations (partially overlapping). Four different situations were tested, as demonstrated on the schematic with the shanks, where the green rectangle indicates the imaging plane. The electrode locations are denoted as ‘A’ and ‘B’. The black arrow indicates the current direction from central electrode A to return electrode B, the red arrow indicates the hypothesized shift of the centroid, compared to base situation ‘A & B in’, where both electrodes A and B are located inside the imaging plane. **(F)** Projection of activated neurons from the 2-dimensional space onto a 1-dimensional axis as defined by the cortical laminar structure to determine the centroid of the population. The projection axis shown here is an axis along the cortical layer. Blue and red circles represent the original and projected neurons, respectively. **(G)** Corresponding Gaussian fit (red curve) of the cumulative ΔF/F of the bins of projected neurons (blue circles). **(H)** Activity centroid location for the four stimulation patterns. The centroid shifts towards the electrode inside the imaging plane, with the red arrow indicating the hypothesized centroid shift and the dashed line representing the midpoint between electrodes A and B. **(I)** The number of activated neurons in the imaging plane decreased when one or more electrodes of the bipolar pair were outside the imaging plane. **(J)** The 3D-simulation model allowed to study the third dimension in the simulations, which included the neurons activated by a current direction out of the imaging plane. **(K)** Horizontal view on the neurons in the 3D-simulated space. The red rectangle defines an area around the stimulation and return electrode, the green rectangle defines the same area but under an angle of 36° with the direction of the stimulation. This represents the angle between the imaging plane and current direction in vivo (panels E-I). **(L)** The number of activated neurons in this rectangle in silico was lower than the number of activated neurons in the rectangle around the bipolar electrode pair, similar to panel I. **(M)** A half sphere depicting the conversion of a possible selectivity increase from a 2D-plane into a 3D-space, with the green dot representing the central electrode, the red dots the variable return electrodes, and the green arrows an example set of possible current directions.

While recordings were limited to the imaged plane, simulations accounted for 3D spatial activation. Applying PCA to the active neurons across L2/3 and L4 for each bipolar electrode pair, we observed a similar number of activation patterns in simulations (5-7 activation patterns, Fig. 5C, D). This consistency across experimental and simulated results underscores the reproducibility of selectivity enhancement with bipolar stimulation.

To further explore the spatial effects of stimulation, we conducted additional recordings where one electrode was positioned outside the imaging plane, creating a 36° current trajectory (Fig. 5E). This configuration activated different subsets of neurons, with the largest population recruited when both electrodes were within the imaging plane (Fig. 5E).

To quantify these activation shifts, we projected the locations of the activated neurons onto the axis between the two bipolar electrodes and calculated the centroid of activation using a Gaussian fit (Fig. 5F, G). The centroid shifted toward the in-plane electrode, likely due to the restricted depth focus of two-photon imaging (Wilcoxon signed-rank test, p = 0.03, Fig. 5H). Neural activity decreased when one or both electrodes were outside the imaging plane (Wilcoxon signed-rank test, p = 0.03, Fig. 5I).

The simulations further supported this effect. When analyzing activation only within a simulated imaging plane tilted relative to the stimulation plane, the number of activated neurons was lower than in the original rectangle area surrounding the bipolar electrode pair (Wilcoxon signed-rank test, p = 0.03, Fig. 5J-L).

The experiment also allowed us to compare the overlap between neural populations activated by two bipolar electrode pairs with two distinct electrodes, in contrast to the previous bipolar electrode pairs where one electrode was always common between different pairs (Fig. 5E). When the current trajectories crossed each other (‘A in, B out’ compared to ‘A out, B in’), overlap was 0.72 (20 µA) and 0.39 (10 µA); when they were parallel (‘A in, B in’ compared to ‘A out, B out’, 100 µm distance between the electrode pairs), overlap was 0.49 (20 µA) and 0.23 (10 µA). This demonstrates a low overlap in neural activity when two different bipolar electrode pairs are located close together, but don’t share an electrode, which can be used to determine the maximum useful density of electrodes in the cortex.

Finally, we estimated the dimensionality of the neural activation patterns in a 3D volume. Within a 2D plane, we identified 5 distinct activation patterns (Fig 5 A-D), corresponding to an angle of 45° between them. By extrapolating from the 2D half-circle arrangement (45° spacing) to a 3D half-sphere, we estimate a theoretical maximum of 17 distinct activation patterns, assuming equidistant current directions (Fig. 5M). However, since each bipolar electrode pair requires two electrodes, this number is halved, leading to a practical resolution increase of 9 distinct activation patterns in 3D. This conversion assumes relatively similar activation patterns in both laminar directions of the visual cortex, in contrast to the highly anisotropic radial direction [42,43].

### The center of neural activity is only slightly affected by stimulation polarity and asymmetry

Steering current in different directions activated distinct neural populations, shifting the center of neural activity (Fig. 3, Fig. S1). However, it remains unclear how the neural activity varies when both electrodes in a bipolar pair are fixed, and stimulation parameters such as polarity and waveform are varied.

To investigate this, we stimulated bipolar pairs with two fixed electrodes with a variable pulse polarity and waveform asymmetry, and compared the neural response. The central electrode was alternated between cathodic-first and anodic-first configurations, and symmetric waveforms were compared with asymmetric waveforms (identical first phase, four times longer second phase). Despite these varying parameters, the activity patterns remained largely consistent across conditions (Fig. 6A), with high correlation coefficients (Fig. 6B).

**Figure 6:**
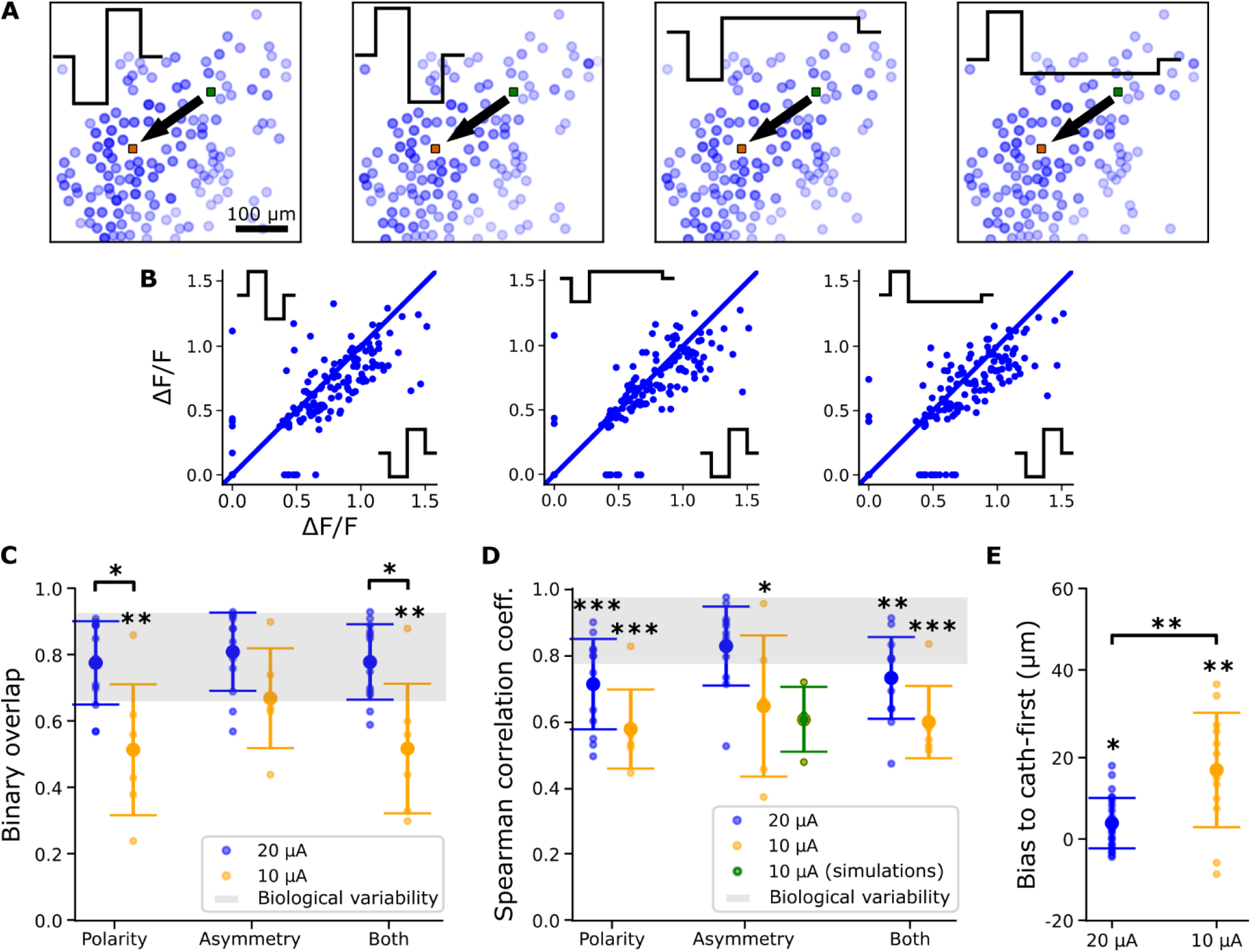
Neural activity was only partially influenced by the polarity and asymmetry of the stimulation waveforms. **(A)** Current sent between fixed central and return electrodes (green and orange squares) with a different polarity and asymmetry activated slightly different neural populations (representative example shown). Four different combinations were tested: cathodic-first and anodic-first, and symmetric and asymmetric waveforms with a factor 4. **(B)** The resulting correlation plots showed a high correlation coefficient: r = 0.82 for variable polarity (left), r = 0.85 for variable asymmetry (middle), and r = 0.83 for simultaneously variable polarity and asymmetry (annotated as ‘both’, right). The identity line is shown in blue, as comparison. **(C)** The average overlap and **(D)** correlation between the neural populations, which was high for all variations of the stimulation parameters. The green datapoints represent simulation results, as comparison. The grey horizontal band denotes the biological variability (mean ± standard deviation), calculated as a 3-by-3 comparison of the repetitions of a certain stimulation pattern in a single slice. **(E)** A small bias of the centroid of activity towards the cathodic-first electrode was noticed, mostly for stimulation with 10 µA.

Polarity variations caused slightly smaller overlaps compared to asymmetry variations, with the influence of both parameters increasing at lower current amplitudes. At 10 µA, the neural overlap was significantly smaller than at 20 µA for both polarity and asymmetry variations (p = 0.01; Fig. 6C). However, no significant differences in correlation coefficients were observed between the two amplitudes (Mann-Whitney U test, p > 0.05, Fig. 6D). These results suggest that while polarity and asymmetry can influence activation patterns, their effects are relatively modest compared to other stimulation parameters such as amplitude or current direction.

To compare these effects with intrinsic biological variability, we conducted a 3-by- 3 cross-comparison of repeated stimulation patterns within a single recording (Fig. 6C, D, grey horizontal band, N=30). The high intrinsic variability among repetitions highlights the high reproducibility of stimulation patterns on the cellular level.

While most measurements approached the observed variability, some differences were still statistically significant. This was the case for the correlation coefficients (variable polarity: p = 0.0009 for 20 µA, p = 0.00007 for 10 µA; variable asymmetry: p = 0.02 for 10 µA; both parameters: p = 0.001 for 20 µA, p = 0.00009 for 10 µA, Fig. 6D), and overlap measurements (variable polarity: p = 0.005 for 10 µA, both parameters: p = 0.006 for 10 µA, Fig. 6C).

To assess the spatial effects of polarity and asymmetry, we computed the centroid of the neural activity by projecting the activated neurons on the axis determined by the two electrodes and applied a Gaussian fit. This centroid of neural activity shifted slightly towards the cathodic-first electrode in bipolar configurations (Fig 6E). This bias was significant at both 20 µA (p = 0.01) and 10 µA (p = 0.003), but it decreased with increasing current amplitude (p = 0.008 for both amplitudes; 1-6 µm 95% confidence interval for 20 µA, 7-26 µm 95% confidence interval for 10 µA). These results align with prior studies indicating greater cortical activation with cathodic-first stimulation, although the impact of the leading-phase polarity decreases for short, high-frequency bursts of multiple biphasic pulses [4,39,40,44–47]. Additionally, it demonstrated that the increased selectivity mostly arose from the location of the bipolar electrodes, rather than the actual polarity of the current direction.

### Bipolar electrode distance can modulate the spatial extent of the neural activation patterns

In addition to enabling selective activation of neural populations, bipolar stimulation offers the advantage of using a local return electrode, which helps limit current spread within the tissue.

To investigate the effect of the distance between the two bipolar electrodes on the neural activation, we compared two configurations: regular inter-shank bipolar stimulation (electrode distance of 150 µm) and intra-shank bipolar stimulation (electrode distance of only 15 µm, located next to each other on the same shank). This resulted in different activated populations, at both 20 µA and 10 µA (Fig 7A, B).

**Figure 7:**
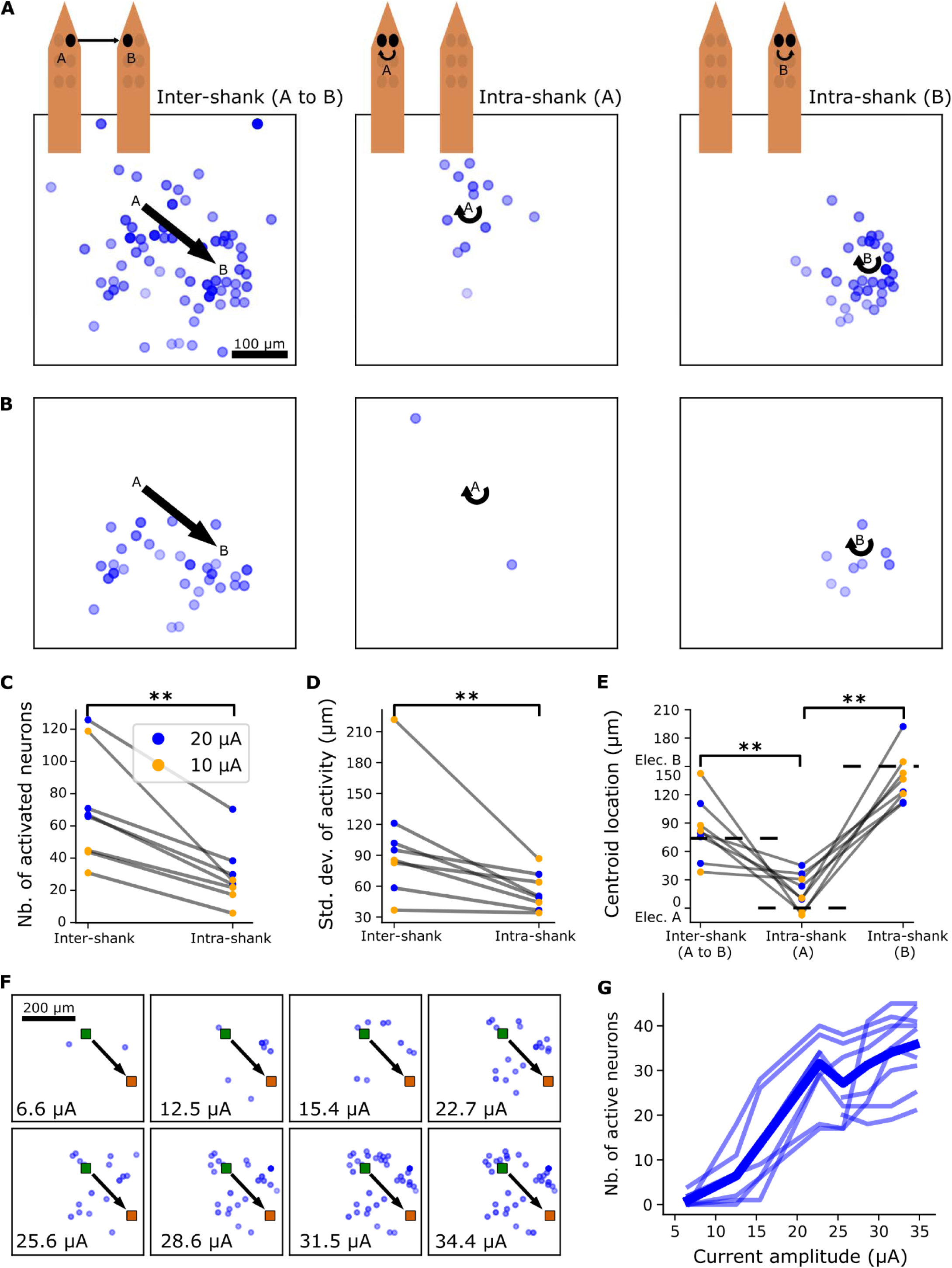
The number of activated neurons and extent of activation was regulated by the bipolar electrode distance and the stimulation current amplitude. Sending current to a return electrode that was only 15µm away (on the same shank) instead of 150µm (on the next adjacent shank) resulted in a much more local neural activation, at **(A)** 20 µA and **(B)** 10 µA. The electrode locations are denoted as ‘A’ and ‘B’. The straight and circular black arrows represent inter-shank and intra-shank stimulation respectively. **(C)** The number of activated neurons and **(D)** extent, quantified by the standard deviation from the Gaussian. Both metrics decreased for intra-shank compared to inter-shank stimulation. **(E)** The centroid of the neural populations for intra-shank and inter-shank bipolar stimulation. The dashed lines represent the hypothesized centroid locations, being at the midpoint between A and B for inter-shank stimulation, and around A and B for intra-shank stimulation at A and B, respectively. **(F)** The number of activated neurons increased with current amplitude, from near-threshold at 6 µA to 36 µA. Fixed central and return electrodes shown as green and orange squares. **(G)** Psychometric curve showing the relation between the number of activated neurons and the increasing stimulation current.

Both the number of activated neurons and the spatial extent of the neural activation patterns were significantly lower when using intra-shank stimulation compared to inter-shank bipolar stimulation (Wilcoxon signed-rank test, p = 0.008 for both metrics, Fig. 7 C, D).

Additionally, the centroid of activity, computed with a Gaussian fit of the projected neurons, shifted depending on the configuration (p = 0.008 for all 3 cases, Fig 7E). For regular inter-shank bipolar stimulation between shank A (0 µm) and shank B (150 µm), the centroid was located between 55 µm and 110 µm (95% confidence interval). For intra-shank stimulation, the centroid was located between 2 µm and 34 µm for shank A, and between 114 µm and 159 µm for shank B (95% confidence intervals).

An increasing electrode distance can be used to increase the number of activated neurons (Fig. 7C). Another stimulation parameter to modulate this, is the current amplitude. We stimulated a bipolar electrode pair with 8 different currents, from near-threshold (6 µA) up to 36 µA (Fig. 7F). The number of activated neurons gradually increased with a higher current amplitude (Fig. 7G). Interestingly, while higher currents typically recruited all neurons activated at lower currents, this inclusion was not always complete, suggesting more complex activation dynamics.

In conclusion, reducing the distance between two bipolar electrodes can be used to decrease the spatial extent of the activation pattern. This demonstrates its use as an extra degree of freedom to finetune stimulation intensity. In the context of visual prostheses, this could enable a more precise control over the brightness and size of a phosphene.

## Discussion

Using custom-designed, flexible intracortical electrodes and two-photon calcium imaging of acute coronal slices of GCaMP6s mice, we investigated the effects of intracortical bipolar stimulation on neural activation. We found that bipolar stimulation enables selective activation of distinct neuronal activation patterns by steering current in different directions. Furthermore, the bipolar electrode distance was found to modulate the spatial extent of the activity.

Unlike prior research in the visual cortex that primarily focused on the centroid shift [13], we directly examined the influence of intracortical bipolar stimulation on single-cell and population-level activation, providing a detailed understanding of how current steering shapes neural activation. This study demonstrates the potential of intracortical bipolar stimulation to enhance neural activation selectivity, significantly exceeding the selectivity achievable with monopolar stimulation. These findings have implications for the development of high-resolution visual prostheses, where precise neural activation will be critical.

### Activation of independent neural populations with current steering

The possibility to activate neurons several hundreds of micrometers away via their axons close to a microelectrode illustrates the importance of the electrical field close to the electrode [9,12,39,46]. Our results highlight the role of the current direction on shaping the activation of neuronal populations.

Most existing examples from current steering or bipolar stimulation take place on a superficial [15,16,19,29–31,33] or a deeper brain area [22,24]. A number of reasons complicate the translation of these prior results to intracortical bipolar stimulation. First of all, the highly non-isotropic laminar structure of the cortex can disrupt the current steering [13,14]. Second, the thresholds and laminar spread of intracortical stimulation in the (visual) cortex are generally depth-dependent [48]. Third, stimulation-induced activity in input layer 4 propagates vertically to layers 2/3 [49]. Despite the impact of the cortical structure on neural activation via intracortical electrical stimulation, we demonstrated variable neural activation both when steering current along a cortical layer (laminar direction), and perpendicular to a cortical layer (radial direction). However, the different structures for the visual cortex of mice and humans require careful clinical interpretation of these results [50,51].

To define the optimal distance between multiple bipolar pairs without showing completely overlapping neural activity, we demonstrated that bipolar pairs with non-overlapping electrodes can be placed at a distance of at least 150 µm and still activate distinct populations (Fig. 5E). This low overlap is in line with earlier research that showed moving an electrode only 15 µm already activated different neuronal populations [39].

Furthermore, we showed that the effects of polarity and asymmetry were less pronounced in altering the neural activation, compared to the current direction.

### Confinement of electrical stimulation using intra-shank return electrodes

Besides current direction, polarity and asymmetry, we also investigated the bipolar electrode distance and amplitude. The decrease in spatial extent of neural activation when using intra-shank bipolar stimulation offers a pathway to minimize unintended neural activation while maintaining efficacy.

In this study, we only implemented intra-shank bipolar electrode pairs with one distance of 15 µm. It is expected that increasing this distance will decrease the confinement of neural activation, and evolve towards regular inter-shank bipolar stimulation. Therefore, the question remains at which distance the bipolar stimulation can be used for current steering and activating different populations, and at which distance the bipolar stimulation acts as confined stimulation with a local return electrode. These are two desired but different effects of bipolar stimulation in the context of a visual prosthesis. Further research should be performed to explore the influence of this intra-shank electrode distance, and how much information it can contribute to the existing set of inter-shank current steering directions, in terms of distinct neural activation.

### Limitations

Despite the promising results, several limitations warrant consideration. First, the two-photon imaging setup was limited to a single plane, which restricts the ability to capture activation patterns in the third dimension. Electrophysiology measurements could provide a detailed view of cellular activity in all dimensions, although these measurements are typically limited to the direct surroundings of electrodes [52–54] and are less suited to study the response to electrical stimulation due to the large electrical stimulation artefacts [55,56]. On the other hand, calcium imaging provides less information about neural activation through electrical stimulation near threshold currents [48].

While the high spatial resolution of calcium imaging allowed us to investigate the effects of current steering on the cellular level, its limited temporal resolution and limited field of view made it impossible to distinguish between direct and indirect activation, and to study the propagation of neural activation to other cortical layers and areas.

While our simulation results support the experimental findings and suggest similar activation patterns in the third unobserved dimension of the cortex [42,57], further experimental validation is necessary.

The simulation environment was based on an existing, previously validated, open-source model from the Allen Institute. We included axons to allow more realistic activation of the neurons in response to extracellular stimulation. The analysis was limited to the layers of stimulation, and therefore neurons relatively close to the electrodes, such that propagation via the axons to different layers was less relevant. Nonetheless, at higher amplitudes or distant locations, the effect of the axons is expected to increase, and the axonal orientation should be further defined and validated [58].

### Future work and clinical implications

In conclusion, this study provides compelling evidence that bipolar electrode configurations and current steering can significantly enhance the precision and selectivity of neural stimulation. By optimizing stimulation parameters such as the current direction and the bipolar electrode distance we could selectively target distinct neural populations and regulate the number of activated neurons. Nonetheless, given the nature of the application, clinical research will be required to investigate how these different neural activation patterns in the primary visual cortex translate to different phosphenes in location, size and brightness, and therefore demonstrating the extent to which current steering and bipolar stimulation can be used to increase the resolution of next-generation visual prostheses.

## Resource availability

Request for further information and resources should be directed to and will be fulfilled by the corresponding author, Vincent Bonin.

## Acknowledgments

The authors would like to thank Laura Apolinário, Michel De Cooman, Thomas Brockhans, Frédérique Ooms, Ine Vlaeminck and Steffen Fieuws for their technical contributions.

## Funding

Vlaams Agentschap voor Innoveren en Ondernemen grant HBC.2021.0187

KU Leuven grant IDN/19/043

HORIZON EIC Pathfinder grant No101071015

## Notes

### Competing Interest Statement

FC is founder of ReVision Implant. MS, LG and FC are employees of ReVision Implant. All other authors declare they have no competing interests.

