## Supplementary Figure 1 for "Intracortical bipolar stimulation allows selective activation of neuronal populations in the cortex"

Maarten Schelles *et al.*

**Fig. S1.**

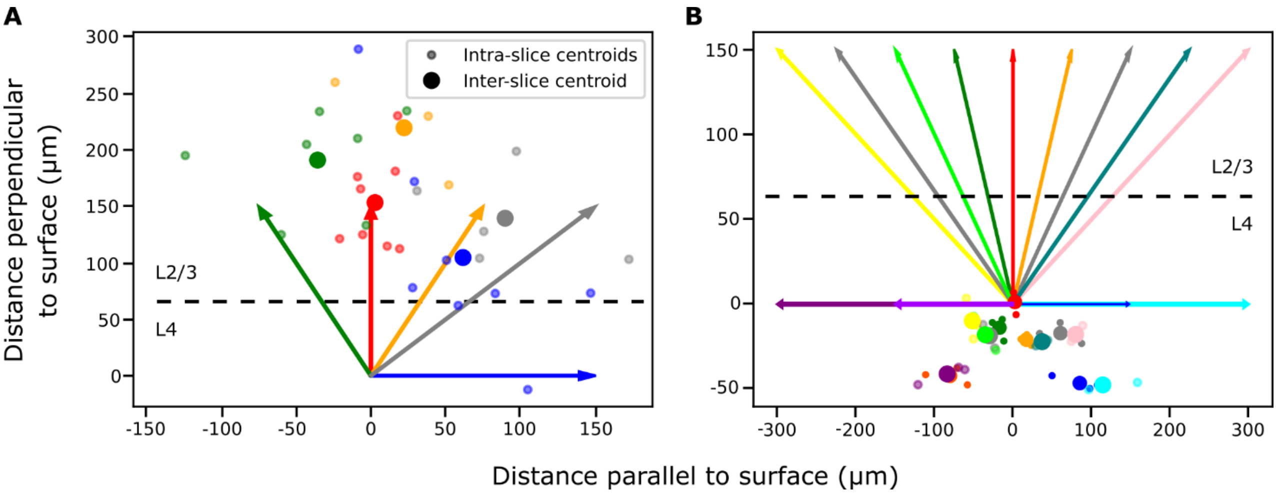


**Supplementary Figure 1: The centroid of the neural activity was influenced by the direction of the stimulation current.** Centroids in a coronal view for a subset of the stimulation directions **(A)** in vivo (20 μA), and **(B)** in silico (10 μA). A dashed line determines the border between L2/3 and L4. The small circles represent the centroids of activity in a certain slice or simulation, the larger circles represent the average of all activity centroids corresponding to a certain direction. Neurons were projected onto two main axes defined by the cortical structure: one parallel to the cortical surface (along cortical layers) and one perpendicular to it. A Gaussian fit on the projected neurons and their amplitude revealed the geometric mean of the activity. The location of the activity in vivo was biased towards L2/3 (A), possibly due to the intrinsically higher number of fluorescent neurons in this layer in this particular animal strain. In contrast, the in silico location was biased towards L4 (B), probably reflecting the higher neuron density in L4 in the model.
